# Structure-based Predictions of Conformational B Cell Epitopes by Protein Language Model and Deep Learning

**DOI:** 10.1101/2025.10.29.685313

**Authors:** Yuhao Zhang, Zhaoqian Su, Felipe Vilicich, Xiaohan Kuang, Yunchao Liu, Grace Zhang, Yinghao Wu

## Abstract

Mapping conformational B-cell epitopes remains a central challenge for antibody discovery: experiments are costly and most computational tools trained on generic protein–protein interfaces transfer poorly to antibody–antigen recognition. We introduce a patch-centric framework that predicts epitopes directly on antigen structures. Each surface “patch” is defined as a triad of neighboring residues, capturing the smallest local unit that encodes both shape and chemistry. We evaluate two classifiers: (i) a protein language model (PLM) approach that averages ESM-2 embeddings over each triad and scores them with a small multilayer perceptron [1], and (ii) a convolutional baseline that consumes a hand-crafted **15*×*20** feature matrix summarizing amino-acid identity, secondary structure, solvent accessibility, and shape index. Trained with five-fold cross-validation on 1,151 AbDb antibody–antigen complexes, the PLM model markedly outperforms the CNN at the patch level (e.g., F1 ***≈* 0.986**, ROC–AUC ***≈* 0.998**). Aggregating patch scores to residues with an ensemble over all folds yields robust residue-wise performance, surpassing the CNN (ROC–AUC **0.689*±*0.072** vs. **0.548*±*0.018**). Against widely used sequence- and structure-based tools on AbDb, our PLM achieves the best summary metrics (ROC–AUC **0.67**, PR– AUC **0.56**) with full coverage of all antigens. On five external complexes unseen during development, the model generalizes well (ROC–AUC **0.663**) and accurately localizes binding regions qualitatively. The method converts PLM representations into interpretable epitope likelihood maps, offering a practical aid for antigen prioritization, antibody engineering, and vaccine design.

## 1 Introduction

Antibodies are produced by B cells in response to antigens, substances that the immune system recognizes as foreign, such as bacteria and viruses, or toxins associated with diseases. The interactions between antibodies and antigens are mediated by residues on their binding interfaces. The residues on the antibodies, mainly situated in the complementarity-determining (CDR) regions of their variable domains [2], are known as paratopes, while the residues on the interface of antigens are called epitopes [3]. Each antibody can recognize a particular group of antigens through the interaction between their corresponding paratopes and epitopes with high specificity and affinity, allowing it to effectively neutralize and eliminate pathogens. While antibodies play essential roles in adaptive immunity by facilitating targeted immune responses and contributing to immunological memory for future infections, mapping B cell epitopes on pathogens can enhance our understanding of how antibodies recognize and respond to these proteins. Moreover, with the knowledge of epitopes on particular proteins that cause autoimmune or infectious diseases, therapeutic or diagnostic antibodies can be more effectively engineered to target these antigens, which can largely reduce the risk of side effects relative to traditional small-molecule drugs known for binding to multiple targets. Additionally, identifying epitopes also helps in designing effective vaccines by pinpointing specific regions of pathogens that elicit strong antibody responses. Therefore, research about mapping B cell epitopes on potential antigens can advance the development of novel disease therapies as well as our knowledge of humoral immunity.

Conventional experimental methods have been the cornerstone of epitope mapping for discovering therapeutic antibodies. These approaches include techniques such as immunization and directed evolution that rely on phage or yeast display systems. The emergence of genome sequencing further allowed the discovery of new antigens directly from genomic information [4]. However, these methods are not only time-consuming, taking months to generate and optimize antibodies, but also labor-intensive, involving complex steps such as animal immunizations and extensive screening processes. Second, the diversity of antibodies generated through immunization is restricted to the natural immune response, which may not always yield high-affinity or broadly neutralizing antibodies. Moreover, these approaches have been much less successful and efficient in targeting those epitopes that are formed by the three-dimensional (3D) folding of proteins, known as conformational epitopes. Over 90 percent of the B-cell epitopes belong to this category [5, 3]. In these cases, low-throughput experiments such as X-ray crystallography or nuclear magnetic resonance may be ultimately required to study the interactions between antibodies and their targeted antigens.

Computational methods offer an ideal alternative for testing conditions that are currently inconvenient to perform in the laboratory. Molecular dynamics (MD), protein-protein docking, and homology-based modeling, or a combination of these algorithms, are the most widely used methods for modeling antibody-antigen interactions [6, 7]. A significant limitation of these methods originates from the fact that the CDR regions of antibodies possess high sequence variations and structural flexibility [8]. As a result, good structural templates for CDR conformations are not sufficient in the protein data bank (PDB). For the same reason, the current force fields for antibody-antigen docking are not accurate enough. The new advances in machine learning (ML), especially deep learning, to study protein-protein interactions (PPIs) have gained enormous attention. For instance, AlphaFold has become highly successful in predicting the structures of PPIs [9, 10]. However, almost all these methods were trained on more generalized datasets of PPIs. They are not sensitive enough to recognize the specific interactions between antibodies and antigens. AlphaFold-Multimer substantially improved general protein complex modeling, but independent benchmarks report limited success on antibody–antigen complexes owing to CDR-H3 variability; early analyses of AlphaFold 3 show improvements yet highlight persistent weaknesses for antibodies [11, 12, 13]. Relative to these generic PPIs prediction methods, a long-standing but still unsolved question in the modeling of antibody-antigen interaction is whether we can predict the conformational epitopes in a given structure of antigen. Various machine learning-based models have been applied to this problem [14, 15]. Unfortunately, a recent study showed that most current methods can only achieve performances that are slightly better than random prediction [14, 15]. Therefore, the development of new deep learning methods that specifically focus on the prediction of conformational epitopes is highly demanding.

This paper delivers a new computational method to predict the potential epitopes given a protein structure. We designed a new representation, which we call local structural patches, to decompose the structure of an antigen into small conformational pieces. Leveraging the recent development of natural language processing, the protein language models (PLMs) showed advantages in predicting a large variety of protein functions [16, 1]. We thus combine a PLM with a a convolutional neural network (CNN) to infer whether a three-residue surface patch belongs to an antibody-binding epitope, using this triad as the smallest unit sufficient to encode local shape and chemical environment. We also compared it with a baseline model that uses traditional structural features. A recently developed large-scale structure-based database of antibody-antigen complexes has further been utilized to train the model. Our cross-validation results show that the PLM model can reach an overall accuracy of about 70 percent, which is much higher than the baseline model. It also outperforms all the other state-of-the-art computational methods using the same benchmark data set. We further applied our model to an independent test set, and our predicted conformational epitopes showed a strong correlation with the experimentally measured data. Together, this study can compensate for the limitations in current experimental or computational techniques for antibody development and could provide potential insights into vaccine design.

## 2 Methods

### 2.1 Structure datasets

In our study, we utilize a curated collection of non-redundant antibody–antigen complexes from the AbDb database [17]. Complexes are clustered by aligning the Fv region sequences of the antibodies; within each cluster, shorter sequences are discarded and the structure with the highest resolution is selected as the representative. Additionally, we exclude complexes in which the antigen contains fewer than 50 residues. This quality-control process results in a refined dataset comprising 1151 antibody–antigen structures.

### 2.2 Surface residues, epitope residues, and local structural patches

For each antigen in an antibody–antigen complex, PyMOL is used to compute the relative solvent-accessible surface area (SaSaRate) of each antigen residue in the absence of the antibody [18]. Residues with a SaSaRate above 0.1 are defined as surface residues. A surface residue is then considered an epitope residue if any of its non-hydrogen atoms lies within 4.5 Å of any non-hydrogen atom of the antibody [19].

We define a local structural patch as any group of three surface residues whose sidechain centers of mass are all within 8.5 Å of one another. A patch is labeled *positive* if all three surface residues are epitope residues, and *negative* if all three are non-epitope surface residues.

### 2.3 Local Structural Patches and Feature Representation

In our **PLM-based model**, we use Evolutionary Scale Modeling (ESM-2) embeddings for each residue within a local patch. ESM-2 is a deep protein language model built on a transformer architecture, trained via masked language modeling on large-scale protein sequence data to capture both local and global contextual signals. For each patch, we aggregate the 320-dimensional ESM-2 embeddings of the three residues by averaging them into a single 320-dimensional vector. This vector then serves as input to a fully connected network that classifies the patch as either a positive (epitope) or negative (non-epitope) patch.

In our **baseline approach**, each patch is transformed into a two-dimensional matrix encoding physicochemical and structural information. First, we calculate solvent accessibility (SaSaRate) and secondary structure (SS) via PyMOL. The shape index (SI) is then derived from an in-house mesh-based curvature analysis, where we convert the antigen’s PDB file into an.obj format to capture its 3D geometry. A triangulated surface mesh is generated from this.obj file, and discrete Gaussian curvature (*K*) and mean curvature (*H*) are estimated at each vertex. The principal curvatures *κ*_1,2_ are subsequently obtained by

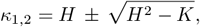

and the shape index *S* is then defined as

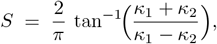

with highly concave regions yielding values near −1 and highly convex regions near +1 [20]. By mapping each PDB atom to its nearest vertex in the mesh, we store the shape index of the corresponding vertex in the atom’s B-factor field.

Next, for each residue in a patch, we apply one-hot encodings for amino acid type (20 classes), secondary structure (3 classes), and solvent accessibility (5 discrete bins). We then combine SS and SaSaRate encodings into a 15-dimensional vector, while amino acid type information selects the appropriate column in a 15×20 matrix, further scaled by the residue’s interaction strength indicator (SI). Contributions from all three residues are summed to produce a single 15×20 matrix that captures local physicochemical composition, structural context, and interaction potential. Finally, these matrices are fed into a convolutional neural network (CNN) to classify each patch as epitope (positive) or non-epitope (negative).

### 2.4 Convolutional Neural Network for Patch Prediction

We trained a convolutional neural network (CNN) to distinguish between positive (epitope) and negative (non-epitope) patches. Each patch was first represented as a 15×20 two-dimensional matrix capturing amino acid identity, solvent accessibility, secondary structure, and shape index features (Section 2.3). The CNN architecture consisted of two sequential convolutional blocks. The first block applies a 3×3 convolution (16 filters) with ReLU activation, followed by batch normalization and a 3×2 max-pooling layer to reduce the spatial dimensions. The second block uses a 2×2 convolution (32 filters), again followed by ReLU, batch normalization, and 2×2 max-pooling. The resulting feature maps are flattened and passed through two fully connected layers (128 and 32 neurons, respectively), both with ReLU activations. A final linear layer with a sigmoid activation outputs the probability that the patch corresponds to an epitope. We trained the model using the binary cross-entropy loss function, optimized with Adam at a learning rate of 10^*−*4^. During each epoch, batches of 500 patch images were sampled randomly from the training set, and early stopping was employed to guard against overfitting.

### 2.5 PLM-Based Patch Classification Using Fully Connected Networks

For our PLM-based model, each patch is represented by the average of three 320-dimensional ESM-2 embeddings (Section 2.3). We then feed this 320-dimensional vector into a fully connected neural network. Concretely, the model begins with a dense layer of 128 neurons with ReLU activation, followed by a dropout layer (rate 0.3) to mitigate overfitting. A second dense layer of 64 neurons (ReLU activation) processes these features before the network outputs a single probability via a sigmoid-activated dense layer. Training is carried out using the Adam optimizer with a binary cross-entropy loss function and a batch size of 32. We reserve 20% of the data for validation and employ early stopping with a patience of 15 epochs, restoring the best-performing weights to prevent overfitting. This framework classifies each patch as epitope (positive) or non-epitope (negative) by capturing both local sequence contexts (through ESM-2 embeddings) and global variation in epitope propensity.

### 2.6 Training and Performance of Patch Prediction Models

We employed a five-fold cross-validation scheme on the set of 1151 antibody–antigen structures. In each fold, 80% of the data (920 structures) were used for training and validation, while the remaining 20% (230 structures) were reserved for testing. The 920 training–validation structures were further split in an 80:20 ratio, yielding a separate validation subset. We trained both the PLM-based and baseline models to classify each patch as either a positive (epitope) or negative (non-epitope) patch, and we report each model’s performance in terms of area under the ROC curve (AUC), precision, recall, and F1-score across all five folds.

### 2.7 Single-Residue Prediction on the Testing Dataset

Using the trained patch prediction models, we identified all surface patches within the 230 structures allocated for testing. Each patch was assigned a predicted score, which was then mapped back to eachresidue forming that patch. This approach provides a residue-level epitope likelihood while maintaining the patch-level prediction context.

For residues belonging to multiple patches, the maximum value of all patch scores is used as the final prediction score, ensuring that the most significant signal can be retained in the local structure. Finally, min-max normalization is performed on the residue level scores within each protein to maintain consistency in the score scale across different residues and proteins.

## 3 Results

### 3.1 Patch-level performance

We evaluated patch classification with five-fold cross-validation over all 1,151 antibody–antigen structures. In each fold, 80% of the structures (920) formed the development pool and 20% (230) were held out for testing; the development pool was further split 80:20 into training and validation for early stopping and model selection. Both classifiers—the PLM-based and the CNN-based models—predict whether a tri-residue surface patch is epitope (positive) or non-epitope (negative). We report accuracy, precision, recall, F1, and ROC–AUC as mean *±* s.d. across folds (Table 1). The PLM classifier consistently and substantially outperforms the CNN baseline (e.g., AUC 0.9985*±*0.0002 vs. 0.9015*±*0.0055; F1 0.9860*±*0.0008 vs. 0.8524*±*0.0101), indicating that sequence embeddings capture local epitope determinants more effectively than the handcrafted structural channels used by the CNN. These results support using the PLM model for downstream residue-level mapping.

**Table 1.** Patch-level performance: mean and standard deviation across folds.

| CNN model | Mean | Std. dev. |
| --- | --- | --- |
| Accuracy | 0.8166 | 0.0103 |
| Precision | 0.9048 | 0.0033 |
| Recall | 0.8059 | 0.0205 |
| F1 Score | 0.8524 | 0.0101 |
| AUC | 0.9015 | 0.0055 |
| PLM | Mean | Std. dev. |
| Accuracy | 0.9817 | 0.0010 |
| Precision | 0.9933 | 0.0010 |
| Recall | 0.9787 | 0.0007 |
| F1 Score | 0.9860 | 0.0008 |
| AUC | 0.9985 | 0.0002 |

### 3.2 Residue-level performance

For residue-wise evaluation, we used an ensemble of *all* fold-trained patch classifiers rather than selecting a single “best” model per fold. On each held-out test structure, every model produced patch-level probabilities; for each surface residue we first aggregated the scores of all patches containing that residue (mean across patches), and then averaged these residue scores across models to obtain an ensemble probability. A binary call (epitope vs. non-epitope) was obtained by thresholding the ensemble probability at a value calibrated on the validation folds.

We report accuracy, precision, recall, F1, and ROC–AUC as mean *±* s.d. across folds (Table 2). As expected, residue-wise classification is more challenging than patch-level prediction, but ensembling substantially reduces variance and the PLM model remains clearly superior to the CNN baseline (e.g., ROC–AUC 0.689*±*0.072 vs. 0.548*±*0.018), supporting its utility for downstream epitope mapping.

**Table 2.** Protein-level performance: mean and standard deviation across folds.

| CNN model | Mean | Std. dev. |
| --- | --- | --- |
| Accuracy | 0.4817 | 0.0249 |
| Precision | 0.1581 | 0.0158 |
| Recall | 0.6358 | 0.0627 |
| F1 Score | 0.2532 | 0.0250 |
| AUC | 0.5481 | 0.0182 |
| PLM | Mean | Std. dev. |
| Accuracy | 0.5545 | 0.0945 |
| Precision | 0.2006 | 0.0402 |
| Recall | 0.7873 | 0.0755 |
| F1 Score | 0.3182 | 0.0546 |
| AUC | 0.6894 | 0.0724 |

### 3.3 Comparison with existing epitope predictors

Table 3 benchmarks our PLM-based approach against widely used baselines, grouped as sequence-based generic, structure-based generic, and structure-based antibody-specific methods. Metrics reported include ROC–AUC, balanced accuracy (BAC), Matthews correlation coefficient (MCC), and PR–AUC; *N*_antigens_ denotes coverage (number of antigens with valid outputs), and *F*_predicted_ is the mean fraction of residues flagged as epitope at each method’s default operating point.

**Table 3.** Comparison of different methods for epitope prediction.

| Method | ROC-AUC | BAC | MCC | PR-AUC | $N_{\text{antigens}}$ | $F_{\text{predicted}}$ |
| --- | --- | --- | --- | --- | --- | --- |
| *Sequence-based generic methods* |  |  |  |  |  |  |
| BepiPred2 [21] | 0.53 | 0.52 | 0.02 | 0.24 | 101/268 | 52% |
| CBTOPE [22] | 0.46 | 0.47 | -0.06 | 0.19 | 229/268 | 38% |
| *Structure-based generic methods* |  |  |  |  |  |  |
| SEPPA3 [23] | 0.53 | 0.51 | 0.00 | 0.23 | 105/268 | 47% |
| DiscoTope2 [5] | 0.58 | 0.53 | 0.06 | 0.26 | 220/268 | 19% |
| ElliPro [24] | 0.56 | 0.53 | 0.04 | 0.23 | 259/268 | 63% |
| EPSVR [25] | 0.53 | 0.52 | 0.03 | 0.26 | 236/268 | 52% |
| BEpro [26] | 0.58 | 0.53 | 0.06 | 0.27 | 235/268 | 12% |
| epitope3D | 0.41 | 0.41 | -0.17 | 0.09 | 221/268 | 3% |
| **PLM** | **0.67** | **0.63** | **0.20** | **0.56** | 1151/1151 | 71.13% |
| *Structure-based antibody-specific methods* |  |  |  |  |  |  |
| EpiPred [19] | 0.50 | 0.50 | -0.01 | 0.35 | 746/1151 | 49% |

Overall, the PLM model achieves the strongest performance across all summary metrics on the AbDb benchmark, with ROC–AUC 0.67, BAC 0.63, MCC 0.20, and PR–AUC 0.56, and provides predictions for all antigens in our set (1151*/*1151). Relative to the best-performing generic structure methods (e.g., BEpro, DiscoTope2), PLM improves ROC–AUC by*~* 0.09 and more than doubles PR–AUC (from *≤* 0.27 to 0.56), indicating a superior precision–recall trade-off under class imbalance. Compared with the antibody-specific baseline (EpiPred), PLM shows larger gains (e.g., +0.17 ROC–AUC and +0.21 PR–AUC) while also increasing coverage.

We note that coverage varies because some tools fail on a subset of structures or impose input constraints (e.g., chain length or formatting). In addition, *F*_predicted_ reflects each method’s built-in calibration and default threshold; hence PR–AUC and ROC–AUC provide threshold-independent comparisons. Together, these results support that leveraging protein language model representations of local surface patches yields more reliable conformational B-cell epitope predictions than traditional sequence- or structure-only baselines.

### 3.4 Case studies on external complexes

To assess generalization beyond the AbDb train/validation/test split, we evaluated the residue-level predictor on five external antibody–antigen complexes (PDB IDs: 7BBJ, 7C6A, 7CWO, 7DR4, 7VYT) not used anywhere in model development. In Fig. 2, antibodies are shown in purple; antigen residues are colored by prediction (red = predicted epitope, blue = predicted non-epitope). Qualitatively, the model localizes binding sites well on 7CWO and 7DR4, with predicted epitope clusters aligning to the observed paratopes. By contrast, 7C6A is more challenging (rarer interface geometry), for which predictions are more diffuse.

**Fig. 1.**
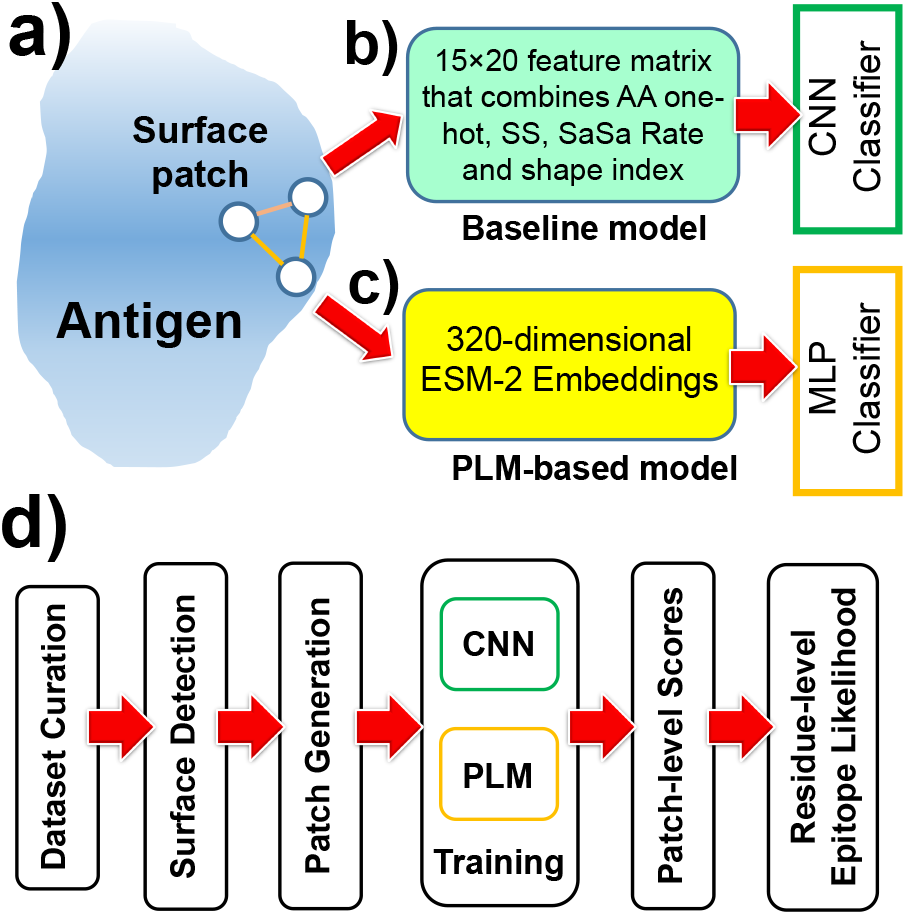
Overview of patch representation and model workflow. **(a)** A surface “patch” is defined as three neighboring surface residues whose side-chain centers of mass fall within 8.5 Å of one another. **(b)** Patch encoding for the CNN: for each residue, amino-acid identity (20-class one-hot), secondary structure (3 classes), solvent accessibility (5 bins), and shape index are combined into a 15 ⨯ 20 matrix, summed over the three residues. **(c)** PLM branch: ESM-2 embeddings (320-D per residue) are averaged across the triad to yield a 320-D vector, then classified by a small MLP. **(d)** Workflow: dataset curation and surface detection → patch generation → training (CNN and PLM branches) → inference with patch-level scores aggregated to residue-level epitope likelihoods.

**Fig. 2.**
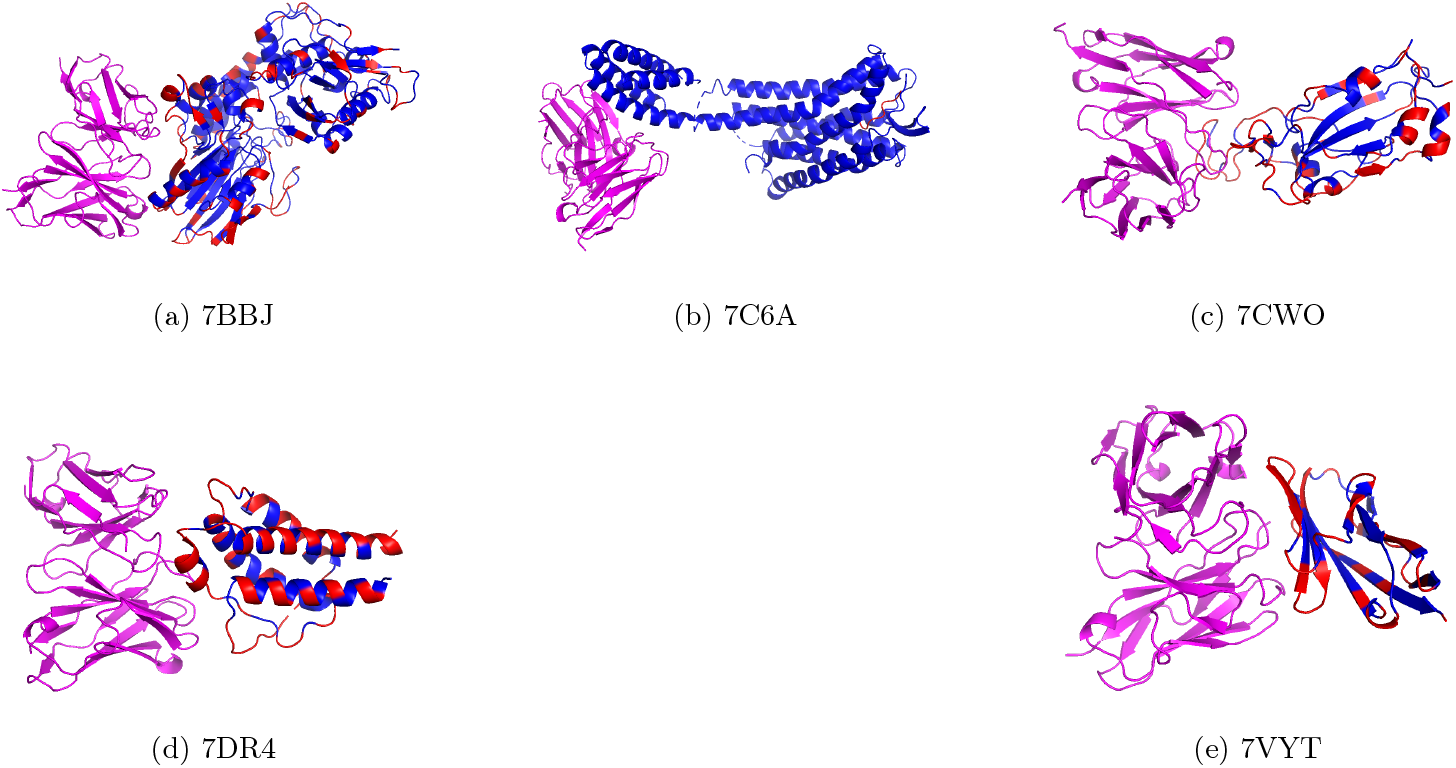
Qualitative results on five external antibody–antigen complexes (not used in training, validation, or testing). Antibody in purple; antigen residues colored by model prediction: red = epitope, blue = non-epitope.

Aggregating over the five cases (Table 4), the model attains ROC–AUC 0.663, with high recall (0.909) but low precision (0.149) at the default threshold—consistent with strong ranking ability but conservative calibration under class imbalance. In practice, the decision threshold can be tuned per application to trade precision for recall.

**Table 4.** Aggregate metrics across the five external cases.

| Metric | Value |
| --- | --- |
| Accuracy | 0.3719 |
| Precision | 0.1489 |
| Recall | 0.9091 |
| F1 score | 0.2559 |
| ROC–AUC | 0.6632 |

## 4. Conclusion

We presented a simple but effective framework for structure-based prediction of conformational B-cell epitopes that operates on local surface patches. By modeling each patch as a triad of neighboring residues, we capture the minimal unit that encodes both local shape and chemical context, and we showed that protein language model features (ESM-2) provide a strong signal for epitope propensity. Across five-fold cross-validation on 1,151 antibody–antigen complexes, the PLM classifier substantially outperforms a CNN baseline built on hand-crafted structural channels at the patch level, and these gains persist when scores are aggregated to residue-wise epitope likelihoods via a simple ensemble. Against widely used sequence- and structure-based predictors on the AbDb benchmark, our approach attains the strongest summary metrics (e.g., ROC–AUC 0.67, PR–AUC 0.56) while producing predictions for all antigens. Qualitative analyses on five external complexes further indicate good localization of binding regions, consistent with the quantitative ROC–AUC of 0.663.

### Practical implications

The model outputs per-residue likelihood maps that can be thresholded for different objectives: high-recall settings are suitable for exploratory scanning of candidate antigens, whereas higher-precision thresholds prioritize focused mutagenesis or paratope-guided design. Because inputs are just antigen structures, the method is easy to deploy upstream of docking or downstream of structure prediction pipelines.

### Limitations and opportunities

One fundamental limitation of current epitope prediction methods lies in the fact that certain regions on a protein surface may be targeted by antibodies that have not yet been experimentally identified, and thus are not labeled as epitopes in existing databases. In other words, current models are trained almost exclusively on experimentally verified epitopes curated in resources such as IEDB [27], leaving potentially immunogenic regions unlabeled. This bias can result in high false-negative rates when predicting novel antibody-binding sites. To address this limitation, future focus could be shifted from predicting known epitopes to predicting intrinsic immunogenic potential. This can be achieved by integrating antibody–antigen docking analysis and molecular dynamics simulations to evaluate the energetic favorability and conformational flexibility of surface patches independent of prior experimental annotations. Such an approach enables the identification of surface patches with high likelihoods of immune recognition, thereby expanding the search space for rational vaccine and therapeutic design.

Our formulation is intentionally local: triads ignore longer-range cooperativity on complex surfaces and do not condition on the cognate antibody. Class imbalance and per-structure calibration also affect precision at default thresholds. Future work will incorporate (i) multi-scale neighborhoods that couple triads with larger geodesic patches, (ii) SE(3)-equivariant graph models to encode richer surface geometry, (iii) antibody-aware conditioning (e.g., predicted paratopes or co-folded complexes) to move toward antibody-specific epitope prediction, and (iv) improved calibration and per-antigen thresholding (e.g., isotonic/Platt scaling) under varying surface compositions.

### Outlook

Despite these limitations, our results show that converting PLM representations into triad-level scores and aggregating them to residue maps yields reliable, interpretable epitope predictions that rival or exceed existing tools. We anticipate this approach will be useful for antigen prioritization, antibody engineering, and vaccine design, and that adding multi-scale geometry and antibody context will further enhance accuracy and utility.

## Author contributions

Z.S. Y.Z., F.V., X.K., Y.L. and Y.W. designed research; Z.S. Y.Z. and F.V. performed research; Z.S., Y.Z.,

F.V. and Y.W. analyzed data; Z.S., Y.Z., G.Z., and Y.W. wrote the paper.

## Acknowledgments

This work was supported by the National Institutes of Health under Grant Numbers R01GM120238 and R01GM122804. The work is also partially supported by a start-up grant from Albert Einstein College of Medicine. Computational support was provided by Albert Einstein College of Medicine High Performance Computing Center. Z.S. thanks the support of the Vanderbilt Data Science Postdoctoral Fellowship. Y.Z., X.K. and Y.L. are grateful for the research funding and support provided by the Vanderbilt Data Science Institute.

